# Detecting color differences in lentil flour samples using an inexpensive, hand-held colorimeter

**DOI:** 10.1101/2024.04.22.590631

**Authors:** Sarahi Viscarra, Ana Vargas, Laura Jardine, Mehmet Tulbek, Kirstin Bett

## Abstract

Our lentil breeding program was interested in determining the effects of the red lentil genotype in processed food products’ colour intensity. This required a colorimeter that could accurately assess colour from the CIE L*a*b* colour space on a small sample of milled flour. Flour colour is commonly assessed with industry standard HunterLab colorimeters that require a large sample size (∼6g flour/sample). In recent years a new spectrophotometer (Nix) has become available. It was designed to quickly and accurately obtain CIEL*a*b* values for everyday items and requires a small amount of sample for analysis (0.5 g flour/sample), but its use has not been tested in milled flour products. To test whether this instrument was appropriate for milled flour colour, eight red lentil genotypes were evaluated for red colour using both instruments. Mean colour scores with Nix (a* and b*) were comparable to those obtained with the HunterLab (r^2^ = **0.9-0.95**), L* scores were less comparable (r^2^ = **0.58-0.61**). Both instruments were able to differentiate and rank red colour intensity in milled flour from the different red lentil genotypes. Nix procedures are more repeatable and use less sample than HunterLab validating its use for assessing red colour intensity in lentil flour.

**Research Highlights:**

- Companies making foods using red lentil flour seek red lentil genotypes that exhibit red colour intensity.
- Nix Spectro2 is an inexpensive photospectrometer made for small volume samples. It has not been tested on flour products.
- The selected red lentil genotypes tested showed a range of red colour intensities.
- Nix Spectro2 performance was comparable to industry standard HunterLab colorimeter.

## Introduction

There is a current demand for alternatives to animal protein for human consumption, and plant-based products have become increasingly desirable. Moreover, pulse-based flours are attractive to industry due to their nutritional value and inexpensive integration to existing product-creation processes. Lentil-based products available to consumers are highly nutritious and meet the demands of specific markets for restricted diets, including gluten-free and low glycemic index products (Campos-Vega et al. 2010). Our program is working with food industry partners interested in developing red lentil-based products like dry pasta, fresh pasta, crackers, and snack foods with a characteristic intense red color in the final product. The partnership was established to determine if certain lentil genotypes or growing environments produced lentil seed with an intense red color that persists through processing from seed to food products. This work required the development of methods to rapidly and cost-effectively assess red color intensity and the persistence of this color intensity through processed products. In similar studies, color has been assessed at different steps of product development with the use of colorimeters or spectrophotometers to obtain scores that represent color through a visual color space (Sant’Anna et al. 2014, Švec et al. 2008, Ugarčić-Hardi et al. 1999). The 1979 Commission Internationale d’Eclairae (CIE) color space (e.g., XYZ, sRGB, LAB) is commonly used to represent color in the form of coordinates that correspond to the changes in the visible space (Becker 2016, Kang 2011). CIE L*a*b* is a three-dimensional uniform color space L* (lightness), a* (green-red axis), and b* (blue-yellow axis) used to describe color and is the best to describe the perception of color by humans, thus it is often used to characterize food (Kang 2011). Colorimetric instruments such as those produced by HunterLab (Reston, VA, USA), among others, are reliable but expensive. This can be a limiting factor for studies constrained by funding availability (Cabas-Lühmann and Manthey 2020).

The food processing industry typically has large amounts of flour samples to work with when assessing color. Getting the desired color in a product is a big focus of their business, and having adequate sample available to get this correct is part of planning for testing. Our food industry partners routinely use HunterLab colorimeters requiring roughly 6 g of sample for flour color testing. Breeding programs do not typically produce large enough volumes of extra seed for these tests so we needed a color assessment method that required less sample than the standard 6g amount used with industry colorimeters.

The Nix Spectro 2 colorimeter (Nix Sensor Ltd., Hamilton ON, Canada), referred to throughout this document as Nix, can be purchased online for one fifth the cost of a standard HunterLab colorimeter. The Nix is a user-friendly device with a mobile app that helps control the spectrophotometer and manage data. From the Nix Toolkit app, settings (e.g., illuminant, observant, technical repetitions, and color space values) can be adjusted as desired, and the scanned values can be directly saved in the app and downloaded later as a .CSV file for further processing. The Nix can also be useful when dealing with smaller sample sizes as it requires roughly 0.5 g of flour/sample. The Nix is routinely used for color assessment of paints, coatings, cosmetics, manufacturing, printing, packaging, and textiles. Recent studies with research applications in food science have demonstrated the Nix can be used to measure color in beef, fish, and lamb (Dang 2021, Holman 2021, Holman 2018). Jha et al. (2021) used the Nix Pro color sensor to assess color in soil and develop prediction models for soil iron content. There are no published studies on the use of Nix for evaluating color in legume or cereal flour. The objective of this study was to validate the use of the Nix as an alternative to a HunterLab colorimeter and describe the methodology applied when determining color in red lentil flour samples.

## Materials and Methods

### Lentil Samples

The red lentil samples used in this study were obtained from the breeding programs at the Crop Development Centre (CDC, University of Saskatchewan, Canada) and at the Università Politecnica delle Marche, Italy. Seeds from genebank accessions were obtained from the International Center for Agricultural Research in Dry Areas (ICARDA) and have been maintained at the CDC over many generations. Three Canadian varieties, one Italian variety, and four genebank accessions were the red lentil genotypes used in this study, and they are described in Table 1. Research plots were grown in Osimo, Ancona, Italy (43°45’04” N, 13°49’98” W), and both Sutherland (52°09’57.3” N, 106°30’16.1” W) and Rosthern (52°44’31.4” N, 106°14’04.5” W), Saskatchewan, Canada, in 2022. Trials in Canada were grown during the typical growing season – from May to August, in plots of 3 m^2^, 3 rows per plot, with inter-row spacing of 30 cm, a row length of 3 m, and a target population of 540 plants per plot. The trial in Italy was grown during February to August in plots of 5 m^2^, 5 rows per plot, with an inter-row spacing of 30 cm, a row length of 3 m, and a target population of 900 plants per plot. Field trials were arranged in a Randomized Complete Block Design (RCBD) with four (2 trials in Canada) and three (1 trial in Italy) repetitions of each lentil genotype for a total of 87 field plots.

**Table 1.** Lentil genotypes evaluated for flour color.

| Accession | Origin | Source |
| --- | --- | --- |
| CDC Redberry | Canada | CDC, USask<sup>a</sup> |
| CDC Robin | Canada | CDC, USask |
| CDC KR-1 | Canada | CDC USask |
| Itaca | Italy | UNIVPM<sup>b</sup> |
| CN 105604 | Ethiopia | PGRC<sup>c</sup> |
| ILL 213 | Afghanistan | ICARDA<sup>d</sup> |
| PI 612875 | Syria | USDA-ARS<sup>e</sup> |
| W 27760 | Egypt | USDA-ARS |

a: Crop Development Centre, University of Saskatchewan, Saskatchewan, Canada, b: Università Politecnica delle Marche, c: Plant Gene Resources of Canada, d: International Centre for Agricultural Research in the Dry Areas, e: United States Department of Agriculture-Agricultural Research Services.

Whole seeds were collected from each plot at maturity and dried at 45°C until a moisture content of <13% was reached. Clean, dried seeds were dehulled using a Satake TM05C huller (Satake Engineering Co. Ltd, Stafford, TX, USA) and finely ground using a Udy cyclone sample mill through a #60 mesh size (0.25 mm) (Udy Corporation, Fort Collins, Co, USA).

### Instrument validation and settings

All measurements were taken in a laboratory under consistent light conditions. The general specifications of both equipment are described in Table 2. The HunterLab Mini XE Plus (HunterLab, Reston, VA, USA), referred throughout this document as HunterLab, was used to obtain the first set of color measurements (Figure 1). Approximately 6 g of each flour sample was placed in a glass Petri dish (75 mm diameter) and measured in triplicate. The HunterLab was standardized using the provided MiniScan XE Plus Reflectance Standard white and black ceramics before every color measurement session. Each slide was placed over the HunterLab’s aperture and scanned (Figure 1, B.2. to B.5). After every scan the generated CIE L*, a*, and b* scores were manually copied to an MS Excel spreadsheet for data analysis and comparison with the Nix results.

**Figure 1.**
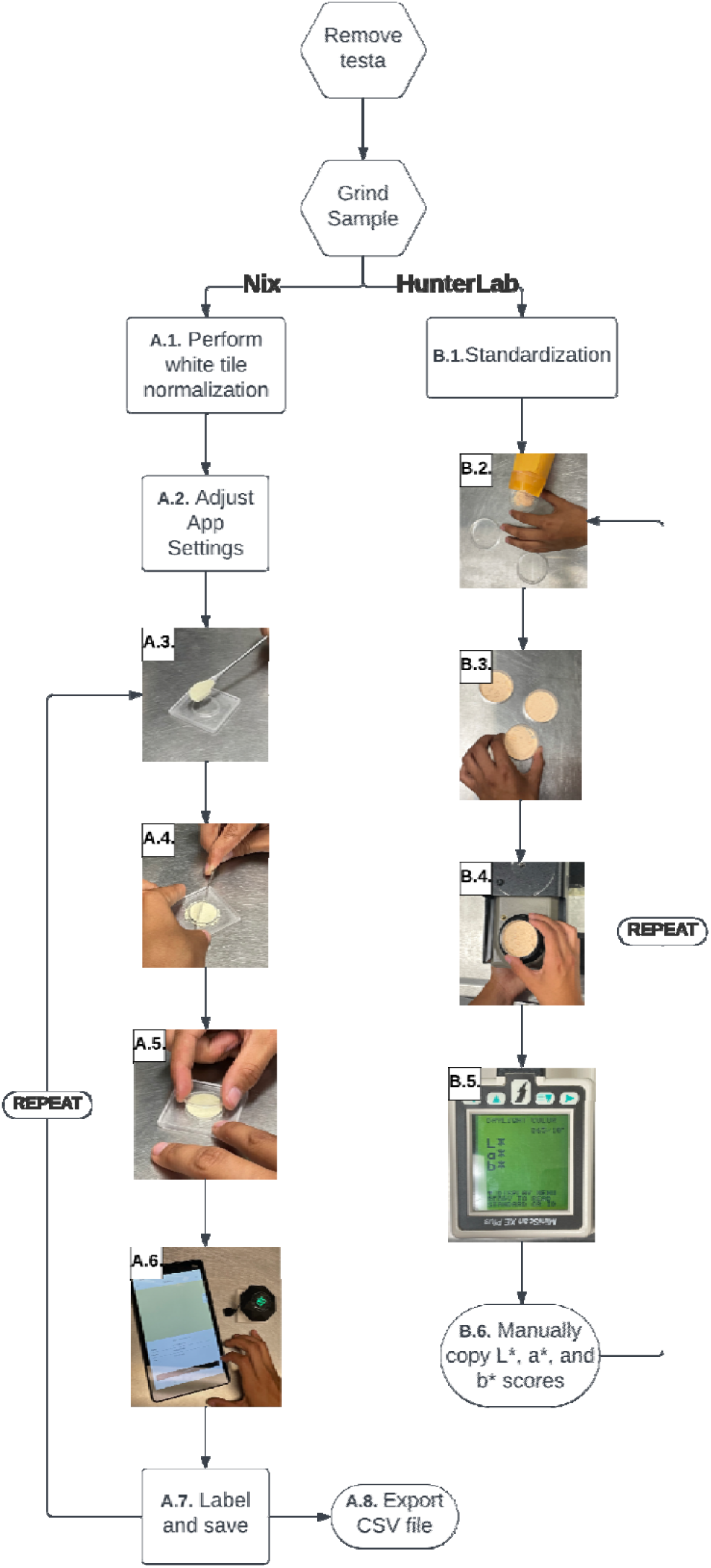
Procedure followed when using the HunterLab MiniScan XE Plus (left) and Nix Spectro 2 (right) for instrument validation. A.1: perform white tile normalization; A.2: adjust app settings A.3: place lentil flour in low-volume adapter; A.4: evenly spread and flatten lentil flour with a small spatula; A.5: place low-volume adapter lid; A.6: scan sample with Nix; A.7: label and save color in library; A.8: Export CSV file to device. B.1: Standardize HunterLab; B.2: place lentil flour in Petri dish; B.3: flatten lentil flour by tapping Petri dish on the table; B.4: place Petri dish on top of HunterLab’s opening and scan; B.5: obtain L*, a*, and b*; B.6: manually copy L*, a*, and b* scores to an Excel Sheet.

**Table 2.**
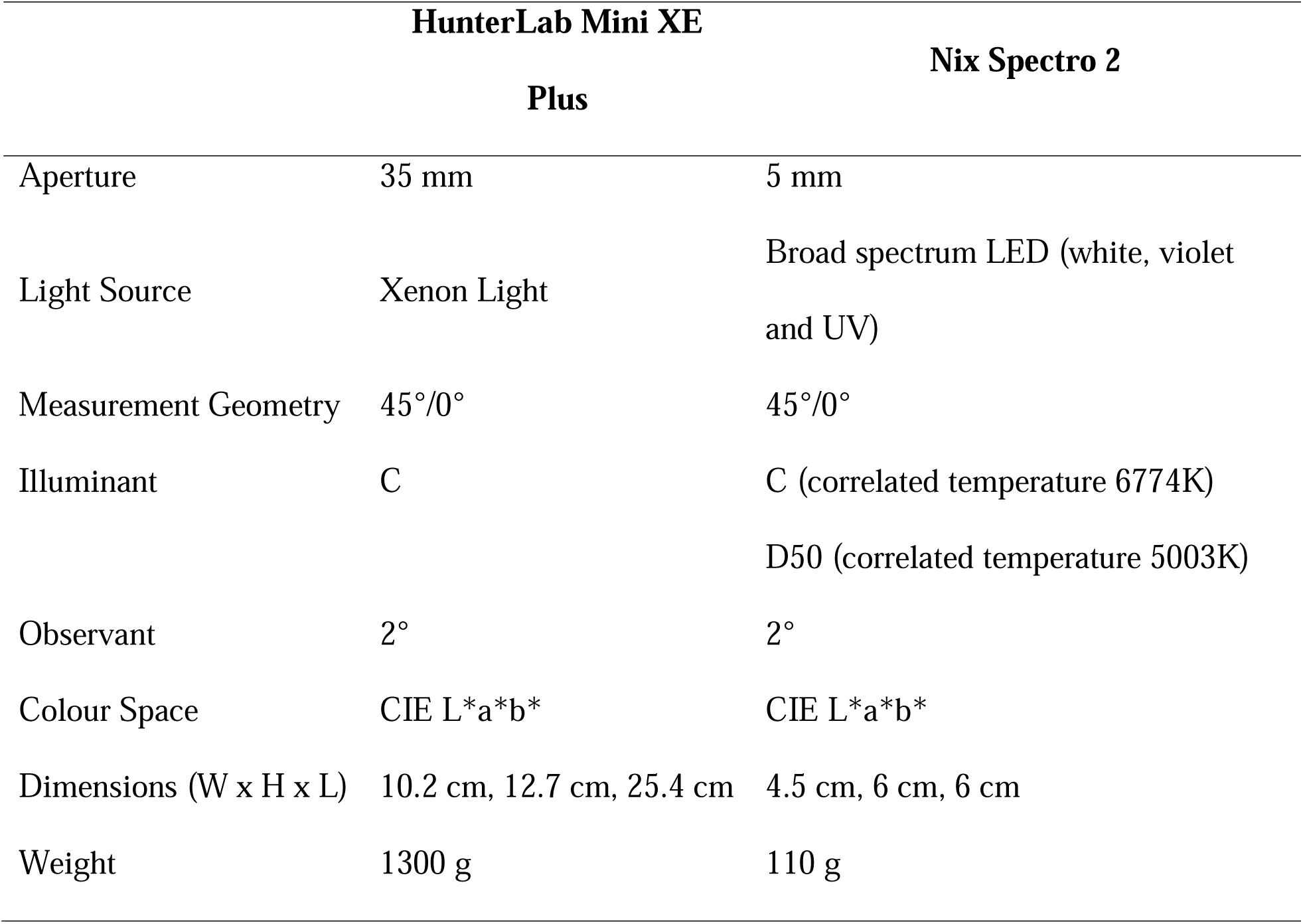
General specifications of the HunterLab MiniScan XE Plus and Nix Spectro 2 used to determine lentil flour color.

|  | **HunterLab Mini XE**<br>**Plus** | **Nix Spectro 2** |
| --- | --- | --- |
| Aperture | 35 mm | 5 mm |
| Light Source | Xenon Light | Broad spectrum LED (white, violet and UV) |
| Measurement Geometry | 45°/0° | 45°/0° |
| Illuminant | C | C (correlated temperature 6774K)<br>D50 (correlated temperature 5003K) |
| Observant | 2° | 2° |
| Colour Space | CIE L*a*b* | CIE L*a*b* |
| Dimensions (W x H x L) | 10.2 cm, 12.7 cm, 25.4 cm | 4.5 cm, 6 cm, 6 cm |
| Weight | 1300 g | 110 g |

A Nix Spectro 2 with illuminants D50 and C was used to scan the same set of samples in triplicate (Figure 1, A.1 to A.7). Illuminant D50 is the default setting of the Nix and is meant to represent natural ‘horizon’ daylight in the early morning or late afternoon, Illuminant C is intended to simulate the average of daylight and commonly used for color measurement in the food and agriculture industry. The data obtained with either can be transformed to the other illuminant values using a tool on the Nix company website: https://www.nixsensor.com/free-color-converter/ if necessary.

For color analysis, the Nix Toolkit App was downloaded on a Samsung Galaxy Tab mobile device and connected to the Nix via Bluetooth®. Every time the Nix was used, a white tile normalization scan was conducted with the provided ceramic reference tile to standardize the spectrophotometer. About 0.5 g of the lentil flour samples were placed in the opening of a clear slide (Nix Pro 2 low-volume adapter, Nix Sensor Ltd., Hamilton, ON, Canada) (Figure 1, A.1 to A.3). The technique allows the same sample to be scanned twice at different points, and the CIE L*, a*, and b* scores obtained from each scan are averaged by the app and saved. This feature is used to accommodate the presence of uneven surfaces or texture. Even though every sample is flattened in the slide during preparation, flour has texture and influences the direction of the light reflected, altering the color values generated – technical repetitions help take these color variations into consideration. Three slides (technical replications) with lentil flour were prepared for every field plot. Each slide was scanned twice, and values were averaged using the multi-point averaging technique in the Nix Toolkit App. The data were collected and stored in a library created in the mobile app and then exported as a .CSV file for data analysis.

### Statistical analysis

Data were analyzed using the general mixed model in R (R Core Team. The R Project for Statistical Computing: https://www.r-project.org/ accessed 30 Oct, 2024). Lentil genotypes were fitted as a fixed effect, and plot repetitions and slides from each plot fitted as random effects. Pearson correlation coefficients between data generated with the HunterLab and Nix were also calculated. Plots were generated using the ggplot2 package in R (Wickham et al. (2016) Create Elegant Data Visualisations Using the Grammar of Graphics: https://ggplot2.tidyverse.org/ accessed 30 Oct, 2024).

## Results

The mean L*, a* and b* values for all eight lentil genotypes varied between the HunterLab and the Nix and between the two Nix illuminants (Table 3). L* scores were the highest with the HunterLab, and the differences between the two instruments were greater than those observed between the two Nix illuminants. The intensity of red color, as measured by a* score values also varied among instruments and showed greatest variation and highest a* score values when samples were evaluated with the Nix using illuminant D50. These were followed by the HunterLab, and the lowest mean was observed with the Nix illuminant C. The b* score values for all lentil genotypes had the lowest variation across instruments, with the HunterLab results having the highest mean. Differences observed among lentil genotypes were significant with both instruments (Table 3, Table S1 for genotype means). The differences observed between means from different field plots were also significant. There were no differences among slide repetitions (technical reps), except for a* and b* score values obtained with the HunterLab instrument.

**Table 3.**
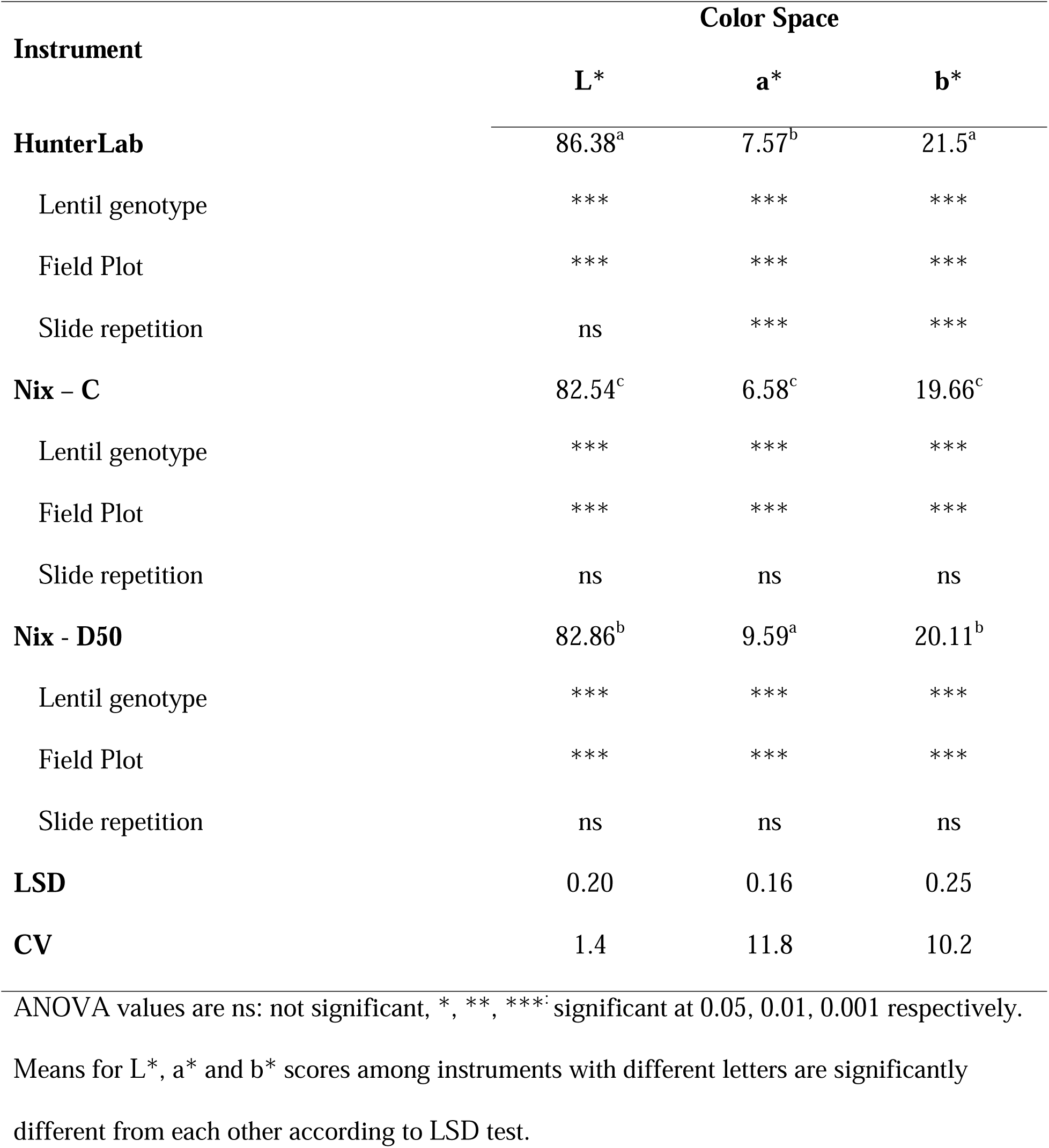
Color space mean values and significance of effects of eight lentil genotypes evaluated with a HunterLab Mini XE Plus and a Nix Spectro 2 with C and D50 illuminants.

| Instrument | Color Space |  |  |
| --- | --- | --- | --- |
|  | L* | a* | b* |
| **HunterLab** | 86.38<sup>a</sup> | 7.57<sup>b</sup> | 21.5<sup>a</sup> |
| Lentil genotype | *** | *** | *** |
| Field Plot | *** | *** | *** |
| Slide repetition | ns | *** | *** |
| **Nix – C** | 82.54<sup>c</sup> | 6.58<sup>c</sup> | 19.66<sup>c</sup> |
| Lentil genotype | *** | *** | *** |
| Field Plot | *** | *** | *** |
| Slide repetition | ns | ns | ns |
| **Nix - D50** | 82.86<sup>b</sup> | 9.59<sup>a</sup> | 20.11<sup>b</sup> |
| Lentil genotype | *** | *** | *** |
| Field Plot | *** | *** | *** |
| Slide repetition | ns | ns | ns |
| **LSD** | 0.20 | 0.16 | 0.25 |
| **CV** | 1.4 | 11.8 | 10.2 |

ANOVA values are ns: not significant, \*, \*\*, \*\*\*: significant at 0.05, 0.01, 0.001 respectively.
Means for L\*, a\* and b\* scores among instruments with different letters are significantly different from each other according to LSD test.

Although the absolute values of color space were different between the two instruments, both were able to distinguish color differences among lentil flour from different genotypes, and the scores obtained with both Nix illuminants were positively correlated with those measured by the HunterLab (Figure 2). HunterLab L* scores had positive correlations with both illuminants of the Nix instrument; however, the correlations were stronger for a* and b* scores between the two instruments measurements for both illuminants.

**Figure 2.**
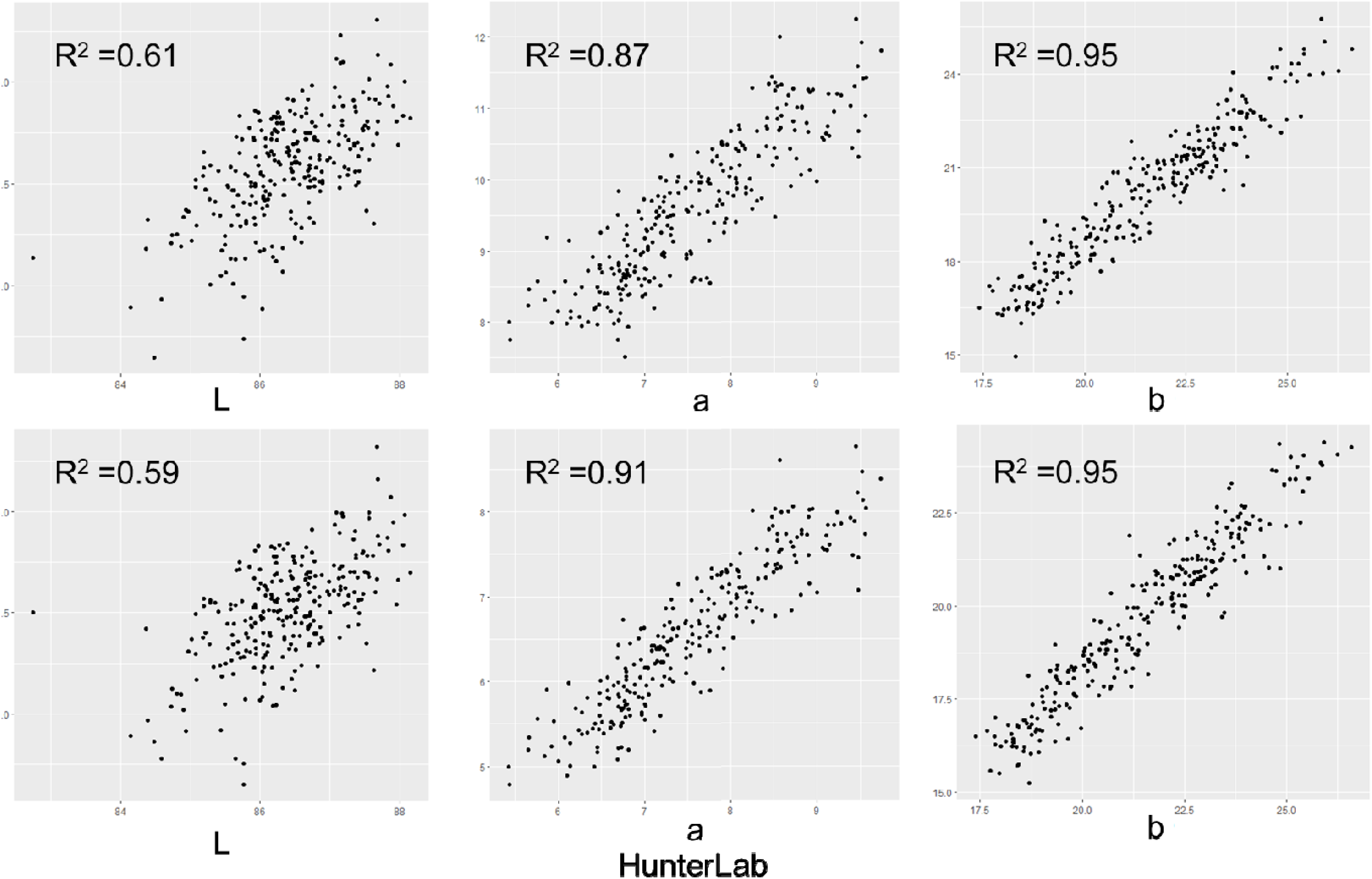
Correlations between L* a* b* color space values of 87 lentil samples evaluated with HunterLab Mini XE Plus and Nix Spectro 2 with C and D50 illuminants.

Differences established among genotypes with the Nix were consistent with those established with the HunterLab (Figure 3, Table S1). Lentil genotypes CDC Robin, ILL 213 and PI 612875, had the lowest L* scores with all three methods, while CDC Redberry, CDC KR-1 and W27760 had the highest values. Similarly, CDC KR-1, ITACA and CDC Redberry, had the lowest b* scores, while CN 105604 and PI 612875 had the highest b* scores, regardless of the colorimeter used. With all three methods the intensity of red color (a*) was higher for CDC Robin and CN 105604, and the highest for PI 612875 among all genotypes with both instruments (Figure 3).

**Figure 3.**
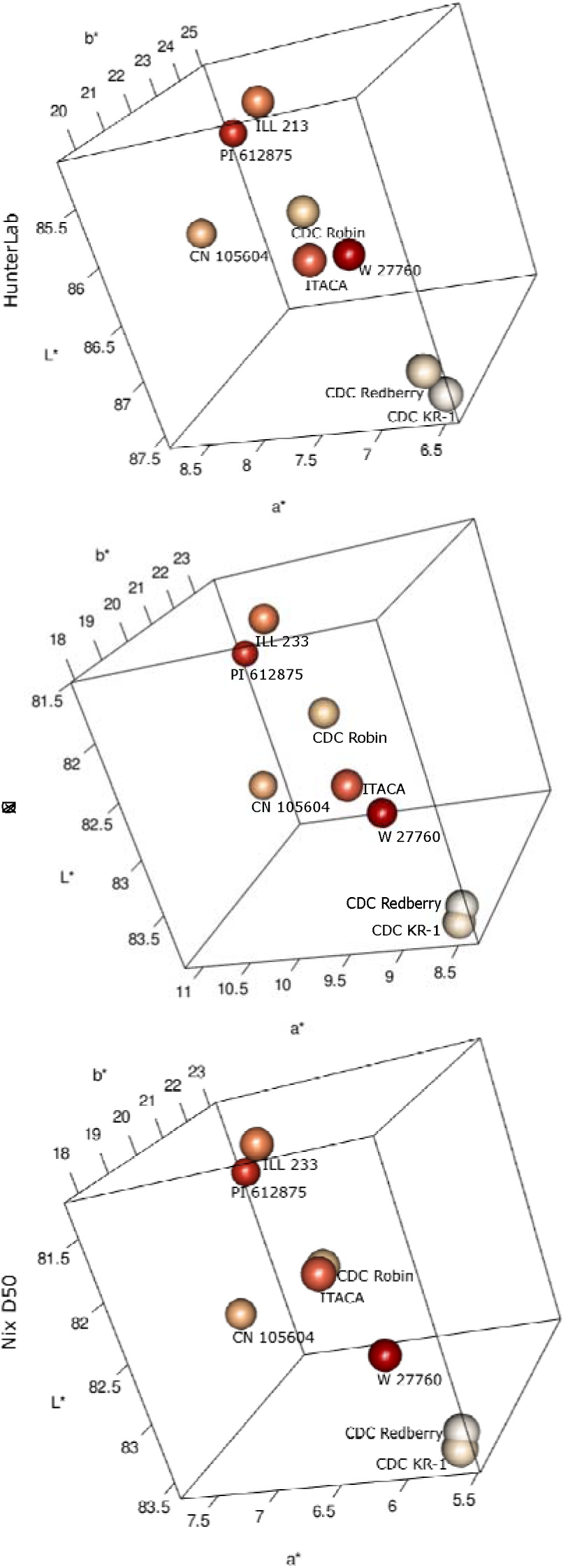
3D plots of the mean L*, a*, and b* scores distribution from the eight genotypes with three different instruments HunterLab (top), Nix illuminant C (middle), Nix illuminant D50 (bottom). L* in the y-axis, a* in the x-axis, and b* in the z-axis.

## Discussion

Color is an important characteristic used to assess product quality (Bicho et al. 2014, Cabas-Lühmann et al. 2020, Sant’Anna et al. 2014, Švec et al. 2008). Breeding programs and industry partners benefit from assessment methods that can be used systematically without incurring time-demanding and expensive phenotyping or requiring a large number of seeds for destructive sampling. The results of this study validated the use of the hand-held, inexpensive Nix instrument and its methodology for assessing color in lentil flour samples in a timely and consistent manner.

Differences in absolute values of L*, a*, and b* scores between instruments and illuminants were expected and have been previously reported (Dang et al. 2021, Holman et al. 2021, Stiglitz et al. 2016, Ugarčić-Hardi et al. 1999). However, the absolute values are irrelevant in this use-case as the main goal was to differentiate color among lentil samples and to accurately identify genotypes with higher a* scores. The genotypes were ordered the same way from the lowest to the highest a* scores regardless of the instrument used and discriminated amongst the genotypes consistently. This validates the use of the Nix as a colorimeter in this application to food color analysis.

The positive correlations observed among L* scores obtained with both instruments were lower than those observed for a* and b* scores. The differences of the L* values found between instruments could be due to differences in methodologies and, or light source differences. It has been reported that temperature and device geometry can influence L* scores (Revantino et al. 2022); however, the same illuminants were used with both instruments, and 45°/0° is the measurement geometry for both (Table 2.) A brighter light source could explain the differences in L* values. While both LED and Xenon Laser are reliable light sources – both cover similar spectral distribution of visible light: 200∼800 nm – LED works best in small spectrophotometers (Gao et al. 2012). The HunterLab uses a Xenon Laser, a brighter light source than broad spectrum LED used by the Nix. Additionally, when scanning samples with the HunterLab the instrument’s aperture is directly exposed to the influence of external light, whereas the Nix aperture is isolated from external light with the help of an adapter. This difference in methodologies can have a direct impact in the L* values obtained. Based in our observations while using both instruments, it is evident that samples have more exposure to external light when using the HunterLab method; another potential advantage of the Nix over the HunterLab for small samples.

It has been proposed that using more slide repetitions for Nix colorimetric analysis would be necessary to improve data precision to account for smaller structural dimensions (Holman et al. 2021, and Jha et al. 2021). We, therefore, expected to have differences among slides (technical reps) due to flour structural variation, but differences were not detected with either illuminant. The flour surface flattening technique used when preparing slides likely helped improve the accuracy of our readings with the Nix, but this difference was not explored in detail during the study. The minor variations observed between technical replicates suggest that fewer repetitions can produce a similar outcome, thus opening up the possibility to scan fewer repetitions per sample and generate the same results in less time. This is particularly useful when sample volume is limited, and data generation time is short. In contrast to the Nix, the HunterLab exhibited differences among slide repetitions for a* and b* scores, indicating that multiple repetitions are needed to obtain more accurate results with this instrument.

The Nix gave us the ability to determine color in much smaller samples than the HunterLab. In breeding programs, phenotyping with destructive methods can be limited by small amounts of seed. Currently, this type of colorimetry is destructive (i.e., requires seed grinding), so at least smaller volumes of flour for measurement is desirable.

Finally, data is more accurately collected and efficiently managed from the Nix Toolkit app. Manually transferring data from the HunterLab to an MS Excel file increases the risk for error or data loss; whereas the automated handling of L*a*b* scores via the Nix App reduces human error and simplifies data management.

The Nix Spectro 2 will be used systematically in our breeding program for determining color in lentil flour of populations developed for different research purposes. In addition to validating the use of the Nix, this work identified genotypes PI 612875 and CN 105604 as the most intense red color lines among evaluated individuals. The same genotypes will be further evaluated for color persistence through to final products.

## Conclusions

In this study, the Nix Spectro 2 was validated for determining L*a*b* scores in lentil flour, as an alternative to more expensive colorimeters that also require more sample for analysis. Nix procedures were more repeatable and rigorous, validating its use for assessing color in lentil flour, especially for small samples, a common limitation within breeding programs.

## Abbreviations

CDC: Crop Development Centre
ICARDA-ILL: International Center for Agricultural Research in Dry Areas-International Legume Lentil
USDA-ARS: United States Department of Agriculture-Agricultural Research Services
CIE: Commission Internationale d’Eclairae
UNIVPM: Università Politecnica delle Marche PGRC, Plant Gene Resources of Canada

## Supplementary Information

Supporting information for this study can be found online in the Supplementary Information section at the end of this article.

---

**Supplemental Table S1.**
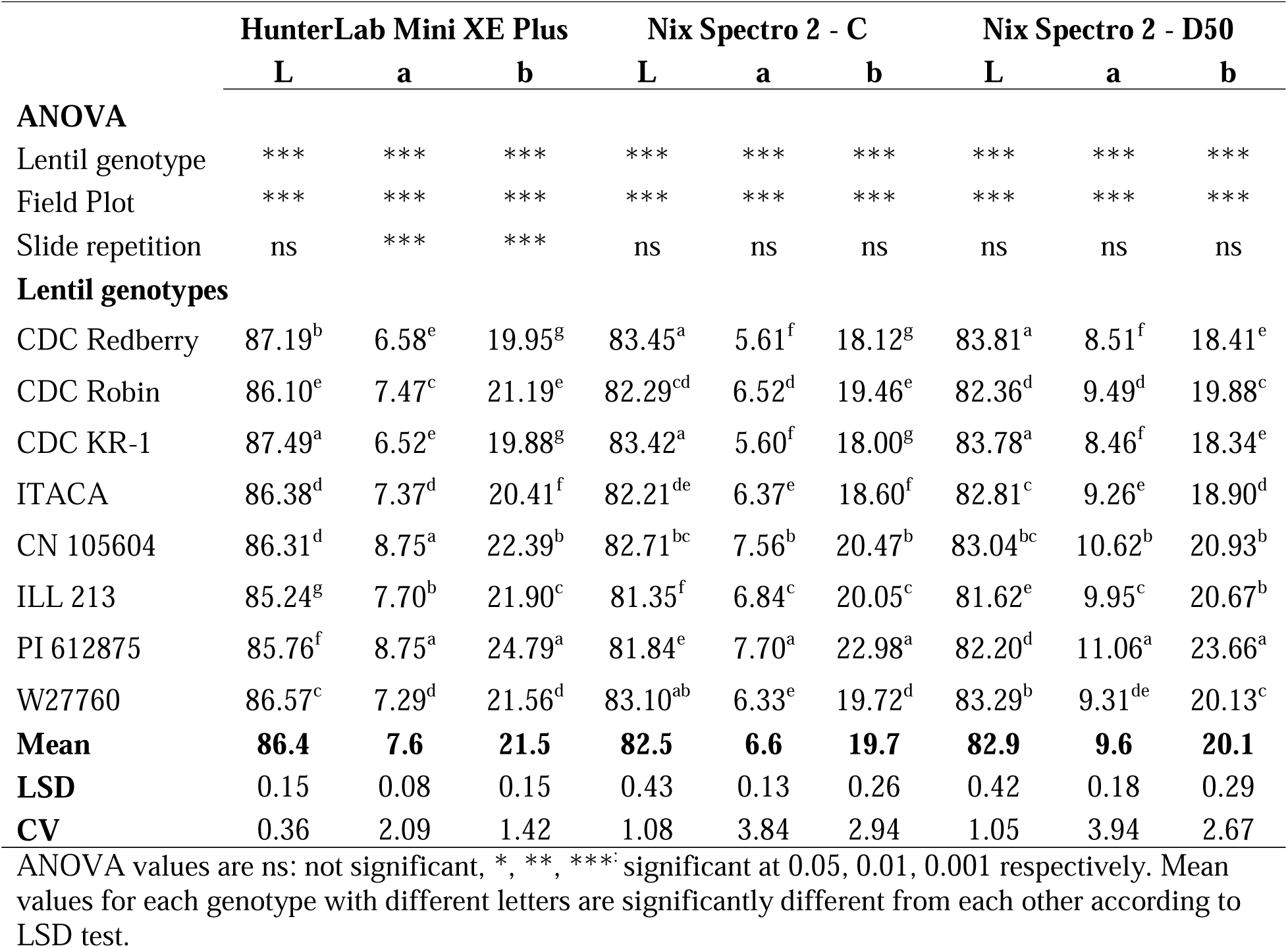
Significance of effects and means for colour space values for eight lentil genotypes evaluated with HunterLab Mini XE Plus and Nix Spectro 2 with C and D50 illuminants.

## Acknowledgements

We thank the Grains Innovation Lab at the University of Saskatchewan for facilitating the use of their equipment and expertise. We also thank the INCREASE team at Università Politecnica delle Marche, Italy for providing samples grown in Italy.

## Declarations

### Ethics approval

Not applicable.

### Consent to participate

Not applicable.

### Consent for publication

Not applicable.

### Availability of data and material

The data supporting this study are available online: https://knowpulse.usask.ca/experiment/LRCS-NixSpectro2

### Code availability

Not applicable.

### Authors’ contributions

Sarahi Viscarra: investigation, data curation, formal analysis, visualization, writing - original draft preparation.

Ana Vargas: conceptualization, investigation, data curation, formal analysis, methodology, visualization, project administration, writing – original draft preparation, funding acquisition. Laura Jardine: conceptualization, methodology, project administration, writing – review & editing, funding acquisition.

Mehmet Tulbek: conceptualization, writing – review & editing, funding acquisition. Kirstin Bett: conceptualization, writing – review & editing, funding acquisition.

